# Interpretable and Generalizable Attention-Based Model for Predicting Drug-Target Interaction Using 3D Structure of Protein Binding Sites: SARS-CoV-2 Case Study and in-Lab Validation

**DOI:** 10.1101/2021.12.07.471693

**Authors:** Mehdi Yazdani-Jahromi, Niloofar Yousefi, Aida Tayebi, Ozlem Ozmen Garibay, Sudipta Seal, Elayaraja Kolanthai, Craig J. Neal

## Abstract

In this study, we introduce and implement an interpretable graph-based deep learning prediction model, which utilizes protein binding sites along with self-attention to learn which protein binding sites interact with a given ligand. Our proposed model enables interpretability by identifying the protein binding sites that contribute the most towards the Drug-Target Interaction. Results on three benchmark datasets show improved performance compared to previous graph-based models. More significantly, unlike previous studies our model performance remains close to the optimal performance when tested with new proteins (ie., high generalizablity). Through multidisciplinary collaboration, we further experimentally evaluate the practical potential of our proposed approach. To achieve this, we first computationally predict binding interaction of some candidate compounds with a target protein, then experimentally validate the binding interactions for these pairs in the laboratory. The high agreement between the computationally-predicted and experimentally-observed (measured) DTIs illustrates the potential of our method as an effective pre-screening tool in drug re-purposing applications.

## Introduction

Drug-target interaction characterizes the binding between a drug and its target, which is critical to discovery of novel drug species, and/or repurposing of existing drugs. High-Throughput Screening remains the most reliable approach to examine the affinity of a drug toward its targets. However, the experimental characterization of every possible compound–protein pair quickly becomes impractical, due to the immense space of chemical compounds, targets and mixtures. This motivates the use of computational approaches for DTI prediction tasks.

Molecular simulation and molecular docking are among earlier computational approaches, which typically require 3D structures of the target proteins to assess the drug-target interaction. The application of these structure-based methods is limited, as there are many proteins with unknown 3D structures, beside that they involve an expensive process. Artificial Intelligence (AI)-based approaches, including Deep Learning (DL) and Machine Learning (ML) algorithms have then emerged to overcome some of these challenges in the process of drug design and discovery. Traditional shallow ML-based models, such as KronRLS Pahikkala et al. [2015] and SimBoost He et al. [2017], require hand-crafted features, which highly affect the performance of the models. Deep Learning has advanced these traditional models due to their ability in automatically capturing useful latent features, leading to highly flexible models with extensive power in identifying, processing and extrapolating complex patterns in molecular data.

Deep Learning models for DTI can be mainly categorized into two classes. One class is designed to work with sequence-based representation input data. Examples of this type include Convolutional Neural Networks (CNNs) and Recurrent Neural Networks (RNNs) that are incapable of capturing structural information of the molecules, leading to degraded predictive power of these models. This motivates the use of a more natural representation of the molecules and the convention of second class of Deep Learning models, namely Graph Neural Networks (GNNs) that use graph descriptions of the molecules, where atoms and chemical bonds correspond to nodes and edges, respectively. Graph Convolutional Neural Network (GCNN) and Graph Attention Network (GAT) are the two widely used GNN-based models in computer-aided drug design and discovery Torng and Altman [2019], Lim et al. [2019], Son and Kim [2021]. All these graph-based models use amino acid sequence representations for proteins, which cannot capture the 3D structural features that are key factors in the prediction of drug-target interactions. On the other hand, obtaining the high-resolution 3D structure of the proteins is a challenging task, beside the fact that proteins contain a large number of atoms requiring a large scale 3D (sparse) matrix to capture the whole structure. To alleviate this issue, an alternative strategy has been adopted wherein the proteins are represented by a 2D contact (or distance) map that shows the interaction of proteins’ residue pairs in the form of a matrix Jiang et al. [2020], Zheng et al. [2020]. It is worth mentioning that a contact (or distance) map is typically the output of protein structure prediction, which is based on heuristics and provides only an approximation abstraction of the real structure of protein, generally, different from the one determined experimentally via X-ray crystallography or by nucleic magnetic resonance spectroscopy (NMR) Tradigo [2013]. Taken all together, and considering the fact that the binding of a protein to many molecules occurs at different binding pockets rather than the whole protein, in this paper, we represent protein pockets as graphs where the key protein residues correspond to the nodes that are connected based on residue proximity. Our model is inspired by the ones developed for text classification in the field of Natural Language Processing (NLP), and is highly explainable due to its self attention mechanism.

### Contribution

Our contribution can be summarized in three parts. First, we use graph representation of protein pockets as the input for target protein. Given the fact that intermolecular interactions between protein and ligand occur in pocket-like regions of the protein, prediction models that utilize the binding sites (pockets) of the proteins are expected to have better generalizability, compared to those relying on certain patterns present in drug molecules or protein sequences. Second, we devise a self-attention mechanism to make the model learn which parts of the protein interact with the ligand, thus complement the black-box nature of deep learning-based methods and enables interpretability, while achieving better DTI prediction performance. Third, we build an end-to-end Graph Convolutional Neural Network (GCNN)-based model, which (1) automatically learns useful embeddings from the graphs of raw molecules and protein pockets, that is, the embeddings are not fixed, but they change according to the context (i.e., sentence) in which they appear and (2) use the learned embeddings similar to the word embeddings, by treating the drug-target complex as a sentence with relational meaning between its biochemical entities a.k.a. protein pockets and drug molecule. This consideration is motivated by the fact that the structure of drug-target complex can be very similar to the structure of a natural language sentence in that the structural and relational information of the entities are keys in understanding the most important information of the sentence. In this regard, each protein pocket or drug is analogous to a word, and each drug-target pair is analogous to a sentence. More specifically, we hypothesize that self-attention bidirectional Long Short-Term Memory (LSTM) mechanism can be used to capture any relationship between binding sites of a given protein and the drug in a sequence, and thus provide a better understanding of their binding relationships. Finally, we conduct in-lab experimental investigations to test the practical potential of our model in prediction and evaluation of compound-target binding interactions in a real world application. Visualization of the aforementioned method can be found in Figure 1. To the best of our knowledge, we are the first to use attention-based bidirectional LSTM networks to perform a relation classification to capture the most important contextual semantic or relational information in a biochemical sequence (i.e. sentence). Each part of our contribution will be described in Section 2.

**Fig. 1.**
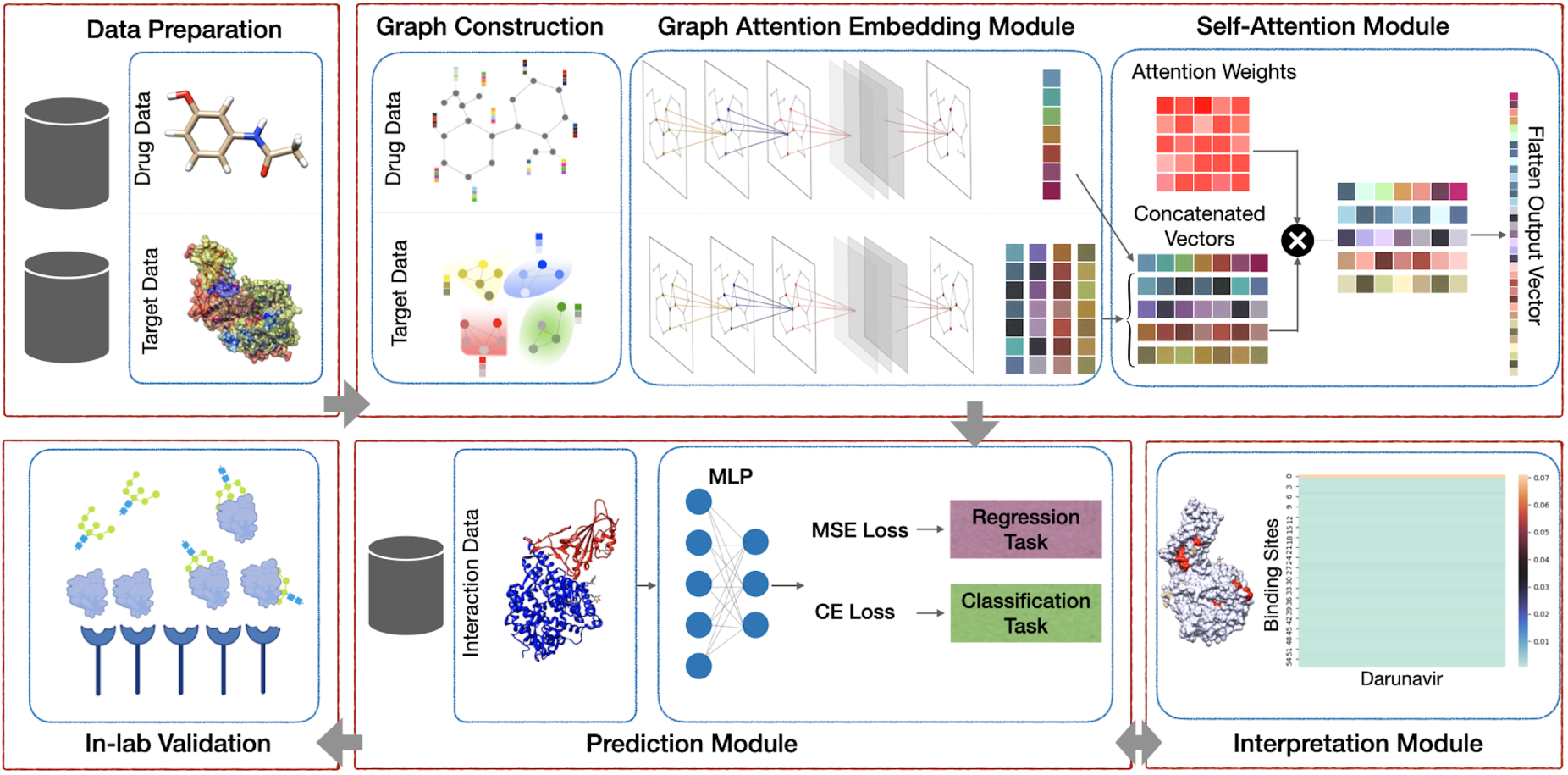
Our proposed framework includes 5 main modules: (1) Preprocessing module that consists of finding the binding sites of proteins; (2) AttentionSiteDTI deep learning module, where we construct graph representations of ligands’ SMILE and proteins’ binding sites, and we create a graph convolutional neural network armed with an attention pooling mechanism to extract learnable embeddings from graphs, as well as a self-attention mechanism to learn relationship between ligands and proteins’ binding sites; (3) Prediction module to predict unknown interaction in a drug-target pair, which can address both classification and regression tasks; (4) Interpretation module to provide a deeper understanding of which binding sites of a target protein are more probable to bind with a given ligand. (5) In-lab validations, where we compare our computationally-predicted results with experimentally-observed (measured) drug-target interactions in laboratory to test and validate the practical potential of our proposed model.

### Related works

Deep learning based approaches have been successfully deployed to address DTI prediction. The main difference between deep learning approaches are in their architecture as well as the representation of the input data. As previously mentioned, small molecules of the drugs can be easily and effectively represented in one-dimensional space, but proteins are much bigger molecules with complex interaction and 1D representations can be insufficient. Although the datasets containing 3D structure of the protein are limited, some recent deep learning based literature have used them in their study. For example, AtomNet Wallach et al. [2015] is the first study that used the 3D structure of protein as input to a 3D convolutional neural network to predict the binding of drug-target pairs using a binary classifier. Ragoza et al. [2017] proposed a CNN scoring function that took the 3D representation of the protein-ligand complex and learned the features critical in binding prediction. This model outperformed AutoDock Vina score in terms of discriminating and ranking the binding poses. Pafnucy Stepniewska-Dziubinska et al. [2018] proposed 3D convolutional neural networks that predicted the binding affinity values for the drug-target pairs. This study represented the input with 3D grid and considers both proteins and ligands atoms similar. Using a regularization technique, their designed network focused on capturing the general properties of interactions between proteins and ligands.

There are limitations associated with all these studies. For example, it is a highly challenging task to experimentally obtain high-quality 3D structure of proteins, which explains why the number of datasets with 3D structure information is very limited Zheng et al. [2020]. Most studies that use 3D structural information utilize convolutional neural networks, which are sensitive to different orientations of the 3D structure, beside the fact that these approaches are computationally expensive.

To overcome these limitations, recent studies have proposed graph convolutional network approaches such as Gomes et al. [2017], Karimi et al. [2019], Nguyen et al. [2021], which take 3D structure of proteins as input for DTI prediction task. There are other studies that applied GCNN to the 3D structure of the protein-ligand complex. Among these studies, GraphBAR Son and Kim [2021], is the first 3D graph convolutional neural network that used a regression approach to predicts drug-target binding affinities. They used graphs to represent the complex of protein-ligand instead of 3D voxelized grid cube. These graphs were in the form of multiple adjacency matrices in which the entries were calculated based on distance and feature matrices of molecular properties of the atoms. Also, they used a docking simulation method to augment additional data to their model. Lim et. al. Lim et al. [2019] proposed a graph convolutional network model along with a distance-aware graph attention mechanism to extract features of the interactions binding pose, directly from 3D structure of drug-target complexes from docking softwares. Their model improved over docking and several deep learning-based models in terms of virtual screening and pose prediction task. However, their approach had limitations such as less explainability as well as addition docking errors added to the deep learning model. Pocket Feature is an unsupervised autoencoder model, which was proposed by Torng et. al. Torng and Altman [2019], to learn representations from binding sites of the target proteins. The model used 3D graph representations for protein pockets along with 2D graph representations for drugs. They trained a GCNN model to extract features from the graphs of protein pockets and drugs’ SMILEs. Their model outperformed 3DCNNRagoza et al. [2017] and docking simlutation models such as AutoDock VinaTrott and Olson [2010], RF-ScoreLiu et al. [2015a], and NNScoreLiu et al. [2015a]. Zheng et al Zheng et al. [2020] pointed out the low efficiency of using direct input of three-dimensional structure and utilized a 2D distance map to represent the proteins. They further converted the problem of drug-target interaction prediction into a classical visual question and answering (VQA) problem, wherein, given a distance map of a protein, the question was whether or not a given drug interacts with the target protein. Although their model outperformed several state-of-the-art models, their VQA system is able to solve a classification task only, where it predicts if there is an interaction between drug-target pairs.

## Materials and Methods

### Preprocessing

We use 3D structure of the proteins that are extracted from Protein Data Bank (PDB) files of proteins. PDB data are collections of submitted experimental values (e.g. from NMR, x-ray diffraction, cryo-electron microscopy) for proteins. We use the algorithm proposed by Saberi Fathi et. al Saberi Fathi and Tuszynski [2014] to find binding pockets of proteins. Figure 2 provides a visualization of a protein’s binding sites. This algorithm computes bounding box coordination for each binding site of a protein. These coordinations were then used to reduce complete protein structures into a subset of peptide fragments. These fragments can be represented as a graph wherein each atom is a node and the connection between atoms are edges in the graph. For each atom, a vector was constructed to represent the atom’s features. Also, one-hot encoding of atom type, atom degree, total number of hydrogen atoms and implicit valence of the atom were used to compute feature vector of each atom. This approach yields vector with a size of 1 × 31 for each node. A bidirectional graph is constructed for each ligand, which is represented in Simplified molecular-input line-entry system(SMILE) format in drug-target interaction data sets. In this study, hydrogen atoms are not explicitly represented as nodes in the graph. Also, a vector was constructed to represent atom’s features in the graph. Similarly, one-hot encoding of atom type, atom degree, formal charge of the atom, number of radical electrons of the atom, the atom’s hybridization, atom’s aromaticity, and the number of total hydrogens of the atom were used to construct the features of the atoms in a ligand. This approach yields vector with a size of 1 × 74 for each node. Generated graphs for proteins and ligands are then fed into a graph convolutional neural network to learn embeddings.

**Fig. 2.**
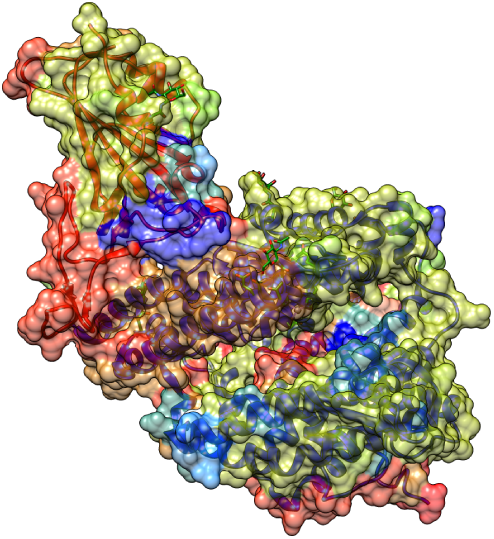
Depiction of COVID spike protein and ACE2 complex with PDB-ID of 6M0J; Color of the surface represents the binding sites computed through Saberi Fathi et. al algorithm which yields the binding site of the proteins. All protein visualization was produced with UCSF Chimera software Pettersen et al. [2004]

### Topology Adaptive Graph Convolutional Networks

We use a Topology Adaptive Graph CNN (TAGCN) Du et al. [2018], which is a variant of graph convolutional and it works by simultaneously sliding a set of fixed-size learnable filters on a given graph. This produces a weighted sum of the filter’s outputs, representing both strength correlation between graph vertices and the vertex features themselves Du et al. [2018]. The graph convolutional layer for TAGCN is defined as

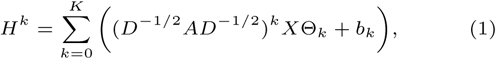

where *A* denotes the adjacency matrix, *D*_*ii*_ = Σ_*j*=0_ *A*_*ij*_ is its corresponding diagonal degree matrix, Θ_*k*_ is the linear weights that accumulates the results of different hops together, with *K* being the number of hops, indicating the length of a path from a given node.

### Pooling Mechanism

Once the constructed graphs for proteins and drugs are fed into a series of graph convolutional layers, we then utilize the method proposed by Li et. al. Li et al. [2017] to extract embedding from the corresponding graphs. For the graph level representation, they define a vector as

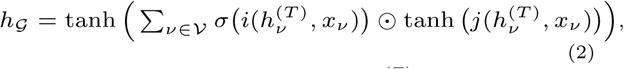

where, *i* and *j* are neural networks, 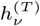 and *x*_*ν*_ are input and outputs real-valued vectors, and 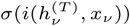 is a soft attention mechanism Li et al. [2017].

### Sequence Handling

Following the extraction of embeddings, we then treat the problem as a text classification problem, which can be defined as follows:

Let *d* ∈ 𝕏 denote a protein-ligand complex, where 𝕏 is space of embeddings for protein pockets and ligands. Also, define the fixed set of classification labels as 𝕔 = {0, 1}, with 0 for non-active and 1 for active interactions for a given drug-target pair. Let 𝔻 denote the labeled training set of protein-ligand complexes ⟨*d, c*⟩, where ⟨*d, c*⟩ ∈ 𝕏 × 𝕔, and it is defined as Eq. (3).

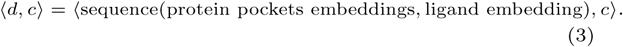

Following the approach proposed by Zhou et. al. Zhou et al. [2016], the goal is to learn a classifier *γ* that maps created sequences to {0, 1}.

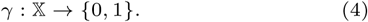

### Self-Attention

Attention mechanism is a method for selectively concentrating on most relevant part of the input vector. It accomplishes this task by mapping a query and a set of key-value pairs to a weighted sum of the values, computed by the relationship of the query and the corresponding key Vaswani et al. [2017]. Vaswani et al. Vaswani et al. [2017] describes a particular attention called “Scaled Dot-Product Attention”, where the input is composed of queries, keys and values. Instead of computing a dot product between the inputs and the query, this attention mechanism contains learnable parameters through adopting three trainable weight matrices. More specifically, as shown in Eq. 5, the queries, keys, and values are packed into matrices Q, K, and V, respectively. Also, the output matrix is computed by calculating the dot product of the query with all the keys, divided by the square root of the dimension of the keys. The division of the square root of the dimension of the keys serves as a scaling factor to avoid pushing the softmax function into small gradient regions Vaswani et al. [2017].

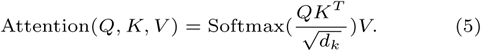

This self-attention mechanism uses sequence of embeddings as input, and extracts query, key and value from each embedding. The attention output is then computed using Eq. 5.

### BiLSTM

Long Short Term Memory (LSTM) is a variation of recurrent neural networks with three gates in its architecture: the input gate, forget gate, and output gate. The cells in an LSTM remember the information in the sequence for an arbitrary index, and the gates regulate the flow of information to and out of each cell. The forget gate, then, decides which information should be forgotten and which information should persist through the next cell. Zhou et al. Zhou et al. [2016] developed a BiLSTM network with two subnetworks for the forward and backward sequence context, respectively. Using an element-wise sum, the outputs of the forward and backward passes are then combined, as shown below:

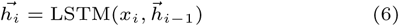

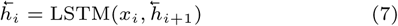

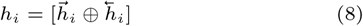

### Classifier

The new values previously computed are then concatenated to a 1D vector *I*, and are passed to the classification layers. In this study, 2 fully connected layers are used to classify concatenated vector to either an active or inactive interaction. Eqs. 9 and 10 represent the two input and output layers of the classifier network, respectively:

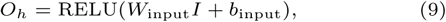

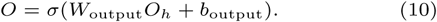

Sigmoid function is then used in the final layer 10 to predict the output in the form of a probability. Moreover, the following cross entropy loss function is used to train the model:

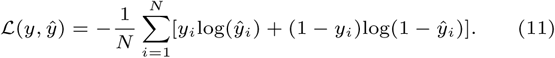

## Experiments

### Datasets

We compare our AttentionSiteDTI with several state-of-the-art methods, using three benchmark datasets: DUD-E dataset, Human dataset and the customized BindingDB dataset. We use the simplest docking-based method to find the binding sites of proteins Saberi Fathi and Tuszynski [2014]. We expect a boost in the performance of our model with incorporation of more complex (ML-based) binding site prediction algorithms, and/or higher-level computational physics approaches (e.g. molecular dynamics, density functional theory).

#### DUD-E

This dataset Mysinger et al. [2012] consists of 102 targets from 8 protein families. Each target has around 224 active compounds and more than 10,000 decoys, which were computationally generated in a way that their physical attributes are similar to active compounds but topologically dissimilar. We used three fold cross validation for our experiment, each fold was splitted based on the target, similar targets were kept in the same fold. We used random under sampling on the decoys to make the dataset balanced for training and used unbalanced dataset for evaluation.

#### Human

This dataset Liu et al. [2015b] was built using a systematic screening framework to create credible and reliable negative sample pairs. the dataset consists of 5,423 interactions. We used the same split used in the DrugVQA Zheng et al. [2020] (80%,10%,10% random split for training,validation and test set) for a head-to-head comparison.

#### BindingDB

This dataset Gilson et al. [2016] contains experimentally based assays of the interactions between small molecules and proteins. Following the work in DrugVQA Zheng et al. [2020], we used a small subset of the dataset, which consists of 39,747 positive and 31,218 negative samples. Further, in order to validate the generalization ability of the proposed model, the testing set was split into two groups of the proteins; those that are seen in the time of training vs those that are not being seen by the model.

### Implementation and evaluation strategy

#### Experimentation strategies

We used Pytorch 1.8.2 (long time support version) for our implementations. We train the models for 30 epochs and used Adam optimizer for training the network with learning rate of 0.001. We used batch size of 100 for better generalization of the network along with a dropout with probability 0.3 after each fully connected layer. The GPU used for the experimentation was (Nvidia RTX 3090) with 24 GB of memory. We used 4 as number of hops in TAGCN for proteins and 2 for ligands. Size of the hidden state for BiLSTM layer in our model was set to 31, which was the output of the graph convolution layer (TAGCN). We used padding of zero to reshape each matrix to the maximum number of binding pockets in the datasets. Also, in order to prevent the attention layer to focus on relationship between different pockets of the protein, the corresponding values for inner protein relationships were set to zero. All other hyperparameters were tuned to yield the best result for each data set, which can be seen in Table 1. *Evaluation metrics*. We evaluated our models in terms of several metrics, including Area Under the receiver operating characteristic Curve (AUC). We additionally report precision and recall for human dataset, accuracy for BindingDB dataset and ROC enrichment metric(RE)for DUD-E dataset, which is defined as

**Table 1.** Dataset’s Training Hyperparameters (FC is representing number of fully connected layers, L-GCN and P-GCN are number of Graph convolutional layers for extracting ligands and protein binding sites embedding respectively)

| Datasets | FC | L-GCN | P-GCN |
| --- | --- | --- | --- |
| DUD-E | 2 | 18 | 16 |
| Human | 6 | 4 | 3 |
| Bindingdb | 6 | 4 | 3 |

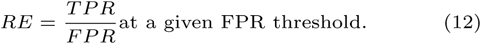

#### Ablation study

Our ablation study illustrates the effectiveness of several text classification methods in AttentionSiteDTI framework. We report AUC of all experiments, which is widely used in the literature. The results of this study can be found in Table 2 showing that the attention mechanism is the most effective method in text classification, and it is particularly advantageous due to its power in explainability. Although, the attention mechanism is showing superior performance compared to Bi-LSTM, it is noteworthy that, for more challenging datasets, attention mechanism cannot capture the relationship between binding sites and ligands. Therefore, for instance in DUD-E dataset, which is intentionally generated in the way that the negative interactions are extremely close to positive ones, attention mechanism with Bi-LSTM architecture gives better results compared to only self-attention mechanism. The Bi-LSTM architecture cannot focus solely on interactions between ligand and binding sites; therefore, it has inferior results compared to other proposed architectures.

**Table 2.** Ablation Study Results

| Ablation tests | BindingDB | Human | DUD-E |
| --- | --- | --- | --- |
| Bi-LSTM | 0.863 | 0.976 | 0.954 |
| Self-Attention | **0.940** | **0.991** | 0.961 |
| Bi-LSTM + Self-Attention | 0.925 | 0.984 | **0.971** |

Finally, the TAGCN architecture for calculating graph embeddings has a better performance compared to GAT Veličković et al. [2018] and GCN Kipf and Welling [2017] architectures, but the performance on other graph convolutional layers is yet to be explored.

#### Comparison on the DUD-E dataset

On DUD-E dataset, we compared our proposed model with several state-of-the-art models that can be divided into 4 categories: (1) machine learning-based methods such as NN-scoreDurrant and McCammon [2011], and Random Forest-score (RF-score)Ballester and Mitchell [2010]; (2) open source molecular docking programs including AutoDock VinaTrott and Olson [2010] and SminaKoes et al. [2013]; (3) deep learning-based models such as AtomNetWallach et al. [2015], 3D-CNNRagoza et al. [2017], which use neural networks to extract features from 3D structural information; and (4) graph-based models like PocketGCNTorng and Altman [2019], GNNTsubaki et al. [2018], DrugVQAZheng et al. [2020], which are all based on graph representations. PocketGCN utilizes two Graph-CNNs that automatically extract features from the graph of protein pockets and ligands to capture protein-ligand binding interactions. CPI-GNNWang et al. [2020] is a prediction model that combines a graph neural network (GNN) for compounds and a convolutional neural network (CNN) for targets. DrugVQA utilizes a 2D distance map to represent proteins in a Visual Question Answering system, where the images are the distance maps of the proteins, the questions are the SMILES of the drugs, and the answers are whether the drug-target pair will interact. Note that the scores of these models are derived form Zheng et. al.Zheng et al. [2020]. Also, following Zheng et al.’s work, we perform 3-fold cross-validation on this dataset, and report the average evaluation metrics. Also, we employ F1 score and ROC enrichment (RE) at thresholds 0.5%, 1%, 2%, and 5%. The results in Table 3 indicate that our model achieves state-of-the-art performance in DTI prediction on all metrics with significant improvement at 0.5% RE. Also, we hypothesize that the poor performance of AtomNet and 3D-CNN may be due to the sparsity of 3D space, as they use the whole 3D structure of the proteins.

**Table 3.** DUD-E Dataset Comparision

|  | AUC | 0.5% RE | 1.0% RE | 2.0% RE | 5.0% RE |
| --- | --- | --- | --- | --- | --- |
| NN Score | 0.584 | 4.166 | 2.980 | 2.460 | 1.891 |
| RF-score | 0.622 | 5.628 | 4.274 | 3.499 | 2.678 |
| Vina | 0.716 | 9.139 | 7.321 | 5.811 | 4.444 |
| Smina | 0.696 | - | - | - | - |
| 3D-CNN | 0.868 | 42.559 | 26.655 | 19.363 | 10.710 |
| AtomNet | 0.895 | - | - | - | - |
| PocketGCN | 0.886 | 44.406 | 29.748 | 19.408 | 10.735 |
| GNN | 0.940 | - | - | - | - |
| DrugVQA | **0.972** | 88.17 | 58.71 | 35.06 | **17.39** |
| AttentionSiteDTI | 0.971 | **101.74** | **59.92** | **35.07** | 16.74 |

#### Comparison on the human dataset

On Human dataset, we compared our model against several traditional ML models such as K-Nearest Neighbors (KNN), Random Forest (RF), L2-logistic (L2)(these results were gathered from Liu et al. [2015a]); and some recently developed graph-based approaches including Graph CNNs(GCNs)Kipf and Welling [2017], CPI–GNNWang et al. [2020], DrugVQAZheng et al. [2020], TransformerCPIChen et al. [2020] as well as GraphDTANguyen et al. [2021] that was originally designed for regression task, and was tailored to binary classification task by Wu et al. [2021]. For a head-to-head comparison with other models, we followed the same experimental setting as in Lim et al. [2016], Tsubaki et al. [2018]. Also, we repeated our experiments with three different random seeds, similar to DrugVQAZheng et al. [2020]. The performances of the aforementioned models were obtained from Wu et al. [2021], and are summarized in Table 4. It can be observed that the prediction accuracy of our proposed model is superior than all ML- and GNN-based models; and it achieves competitive performance with DrugVQA in terms of precision and recall. The relatively low performance of ML-based models is indeed in line with our expectation, and is due to their use of low-quality features, unable of capturing complex non-linear relationships in drug-drug interaction. The deep learning models, on the other hand, are very powerful in extracting important features governing the complex interactions in a drug-target pair. On this basis, our model further improves on the accuracy, indicating that the quality of learned information in drug-target interactions is guaranteed by the back propagation of the end-to-end learning of our AttentionSiteDTI.

**Table 4.** Human Dataset Comparision

|  | AUROC | Precision | Recall | F1 Score |
| --- | --- | --- | --- | --- |
| K-NN | 0.86 | 0.798 | 0.927 | 0.858 |
| RF | 0.940 | 0.861 | 0.897 | 0.879 |
| L2 | 0.911 | 0.861 | 0.913 | 0.902 |
| SVM | 0.910 | **0.966** | 0.950 | 0.958 |
| GraphDTA | 0.960 | 0.882 | 0.912 | - |
| GCN | 0.956 | 0.862 | 0.928 | - |
| CPI-GNN | 0.970 | 0.923 | 0.918 | 0.920 |
| E2E/GO | 0.970 | 0.893 | 0.914 | 0.903 |
| DrugVQA | 0.979 | 0.954 | 0.961 | 0.957 |
| BridgeDPI | 0.990 | 0.963 | 0.949 | 0.956 |
| AttentionSiteDTI | **0.991** | 0.951 | **0.975** | **0.963** |

#### Comparison on the BindingDB dataset

On BindingDB dataset, we further compared our model against TiresiasFokoue et al. [2016], DBNWen et al. [2017], CPI-GNN Wang et al. [2020], E2EGao et al. [2018], DrugVQAZheng et al. [2020] and Bridge-DPIWu et al. [2021] as baselines. Tiresias uses similarity measures of drug and target pairs. DBN uses stacked restricted Boltzmann machines with the inputs in the form of extended connectivity fingerprints. As mentioned earlier, CPI-GNN combines a graph neural network (GNN) for compounds and a convolutional neural network (CNN) for targets to capture drug-target interactions. E2E is a GNN-based model that uses LSTM to learn drug-target pair information with Gene Ontology annotations. DrugVQA, as previously mentioned, is a Visual Question Answering system, where the images are the distance maps of the proteins, the questions are the SMILES of the drugs, and the answers are whether the drug-target pair will interact. Finally, BridgeDPI uses convolutional neural networks to obtain embeddings for drugs and proteins, as well as a GNN to learn the associations between proteins/drugs using some hyper-nodes that are connections between proteins/drugs. Note that the scores for all these models are derived from Wu et al. [2021]. Also, following suggestions from previous works, we report the prediction results in terms of AUC and Accuracy (ACC) on the test set, which is divided into a set of unseen protein (the proteins that are not observed in training set) and a set of seen protein (the proteins that are observed in training set). This, indeed, makes the customized BindingDB dataset suitable to assess models’ generalization ability to unknown proteins, which should be the focus in prediction problems (i.e., cold-start problem), as there are a large number of unknown proteins in nature.

As experimental results indicate in Figure 3, all models generally perform well on seen proteins with AUC above 0.9, and ACC exceeding 0.85. However, these models show different and much worse performance on unseen proteins, which reflects the complexity of this more realistic learning scenario. Tiresias is a similarity-based model that uses a set of expert designed similarity measures as the features for proteins and drugs. The poor performance of Tiresias on the unseen proteins is perhaps due to the fact that these handcrafted features are not sufficient in capturing interactions between drug-target pairs, thus resulting in the accuracy even less than 0.5 on unseen proteins. On the other hand, the good performance of deep learning-based models including DBN, CPI-GNN, E2E, DrugVQA, BridgeDPI as well as our AttentionSiteDTI shows the effectiveness of these models in capturing relevant features that are critical in DTI prediction problem. As the results show, our model achieves the best performance with AUC of 0.97 and 0.94 on seen and unseen proteins, respectively. Also, in terms of accuracy, our AttentionSiteDTI outperforms all other models with accuracy reaching 0.89 in unseen proteins. This is an indication that our attention-based bidirectional LSTM network is, indeed, effective in relation classification of drug-target (protein pocket) pairs by learning the deeper interaction rules, governing the relationship between proteins’ binding sites (pockets) and drugs. Also, the seemingly good performance of baselines on seen proteins can be an indication of over-fitting.

**Fig. 3.**
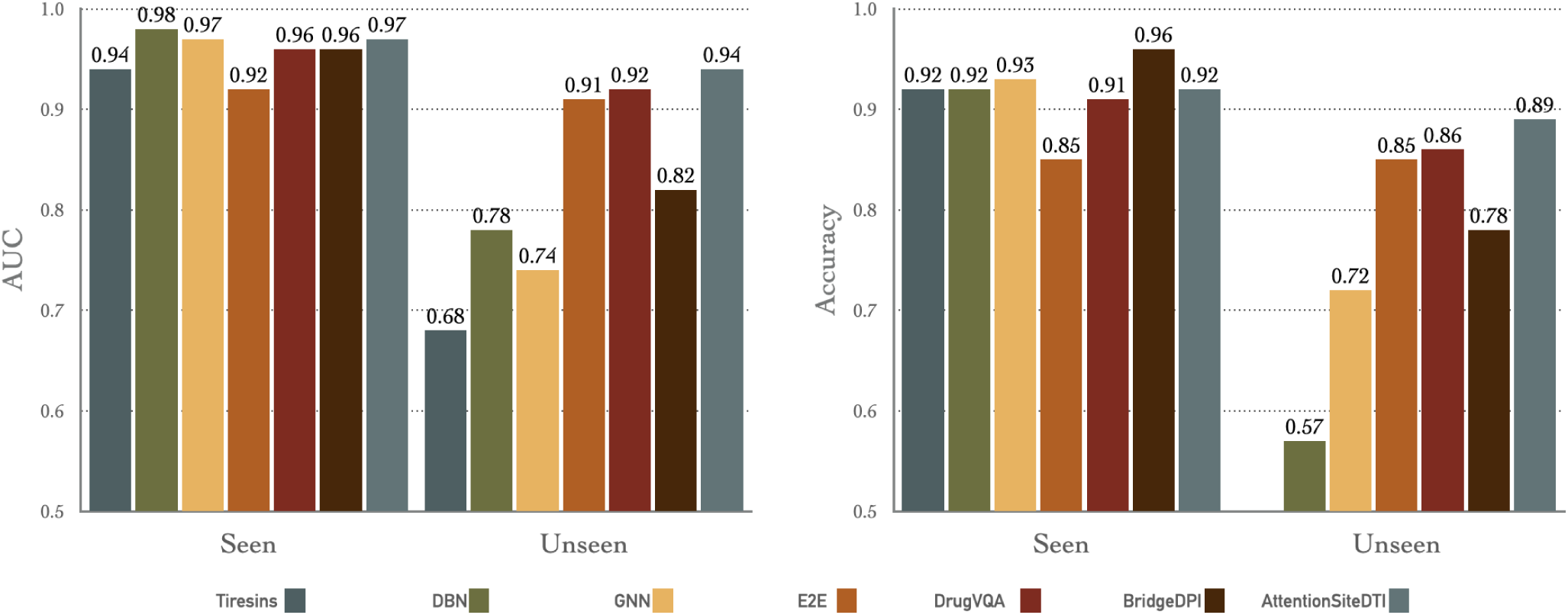
Comparison of AttentionSiteDTI with six baselines: (left) shows Area Under the Curve (AUC) for seen proteins and unseen proteins in the test; (right) shows Accuracy for seen proteins and unseen proteins in the test. Note that the accuracy scores of Tiresias do not show in unseen case, because it is lower than the lower bound of the y-axis (0.5). Note that for a head-to-head comparison with all models including ours, we implemented the BridgeDPI model with our experimental setting. Our model outperforms all other methods in unseen proteins which means our model is better in generalization over other models. In the seen protein scenario our model is comparable to other models and high AUC and accuracy in seen scenario indicates over-fitting of the model.

#### Model Explainability

Ligands bind to certain parts (active sites) of proteins either blocking the binding of other ligands or inducing a change in the protein structure, which produces a therapeutic effect. Binding at other sites that provide no therapeutic value are “non-active,” and generally do not cause a direct biological effect. Ligands binding to active sites and inducing a change in protein structure (conformation) are less likely in our system of study, and are probably not as helpful for building models (usually these ligands/therapeutic agents are employed/considered when a patient has an ailment, which causes natural biochemicals to be produced in insufficient quantities).

In this work the attention mechanism enables the model to predict which protein binding sites are more probable to bind with a given ligand. This probability is the attention matrix computed in the model. The attention visualization can be found in Figure 4 as the heat map of the protein for the complex of SARS-CoV2 Spike protein and human, host cell-expressing ACE2 in the interaction with the drug named Darunavir. The projection of the heat map on the protein is depicted in this figure, as well.

**Fig. 4.**
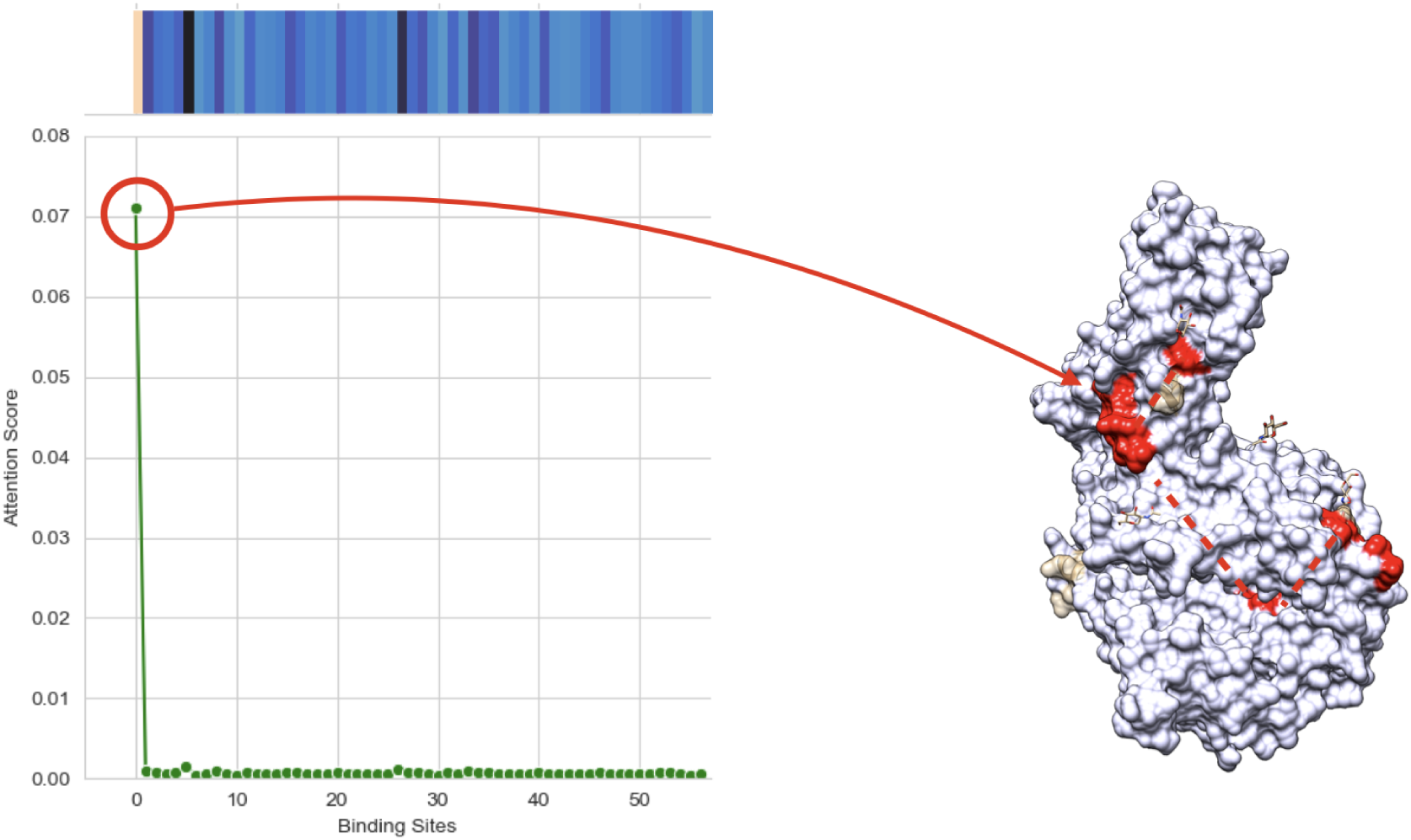
(Left) shows Heatmap and line plot of self-attention mechanism weights for each binding site in the proposed method with input of Darunavir as ligand and complex of COVID spike protein and ACE2 as protein, which translates to probability of each calculated binding sites of the protein being active for that specific ligand.(Right) shows projected heatmap of self-attention weights on the complex of COVID spike protein and ACE2. This figure shows the interpretablity of our model which can give us the binding site that have the most probability of binding to the ligand.

## SARS-CoV-2 Case study and In Lab Validation

To further evaluate the practical potential of our proposed model, we experimentally tested and validated the binding interactions between spike (or ACE2) protein and seven candidate compounds including N-acetyl-neuraminic acid, 3*α*, 6*α*Mannopentaose, N-glycolylneuraminic acid, 2-Keto-3-deoxyoctonate ammonium salt, cytidine5-monophospho-N-acetylneuraminic acid sodium salt and Darunavir as inhibitor molecules to bind to the spike protein, or the ACE2 receptor protein, which has been shown to be the primary host factor recognized and targeted by SARS-CoV-2 Spike protein. The primary goal in our experimental investigations is to determine the ability of those seven compounds to disrupt the important interaction between spike protein and ACE2, which, in turn, leads to inability of SARS-CoV-2 virus to infect host cells. As the results show in Table 5, we observe high agreement (five out of seven matched results) between the predicted and experimentally-measured drug-target interactions, which illustrates the potential of our AttentionSiteDTI as an effective complementary pre-screening tool to accelerate the exploration, and recommendation of lead compounds with desired interaction properties toward their targets. In our experiment, we set the activity threshold to 15 nm to only capture highly active compounds; thereby, limiting the influence of interactions at neighboring sites and weak interactions with poor coordination to the binding site center.

**Table 5.**
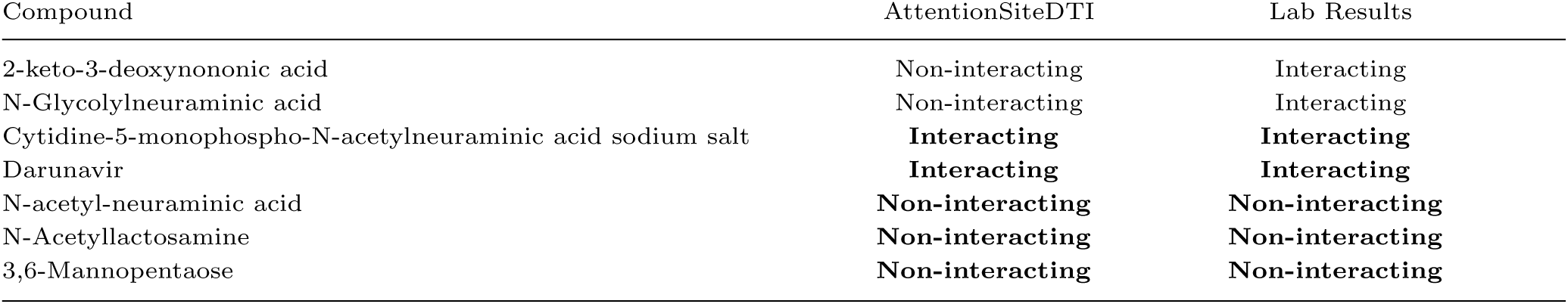
In-lab Validation of AttentionSiteDTI in the case study of Covid-19

## Conclusion

In this work, we proposed an end-to-end Graph Convolutional Neural Network(GCNN)-based model, built on self-attention bidirectional Long Short-Term Memory mechanism, which captures any relationship between binding sites of a given protein and the drug in a sequence analogous to a sentence with relational meaning between its biochemical entities a.k.a. protein pockets and drug molecule. Our proposed framework enables learning which binding sites of a protein interact with a given ligand, thus allows interpretability and better generalizability, while outperforms state-of-the-art methods in prediction of drug-target interaction. We experimentally validate the predicted binding interactions between seven candidate compounds and spike (or ACE2) protein. The results of our in-lab validation showed high agreement between the computationally-predicted and experimentally-observed binding interactions. Our model exhibit state-of-the-art performance, is highly generalizable, provide interpretable outputs and performs well when validated against in-lab experiments. As a result, we expect it to be an effective virtual screening tool for drug discovery.

### Key Points

- We build an end-to-end Graph Convolutional Neural Network (GCNN)-based model for predicting drug-target interactions.
- Self-attention mechanism enables the model to predict the most active binding site of the protein in drug-target interaction which translates to interpretablity of the deep-learning model in this branch of problem.
- The proposed model showed the best generalizability among all other drug-target interaction prediction models.
- We validated our predictions by experimentally testing the candidate compounds.

## Data and Code availability

All datasets are publicly available. DUD-E dataset is available at http://dude.docking.org, Human dataset is available at https://github.com/IBMInterpretableDTIP and finally the customized BindingDB-IBM dataset can be found at https://github.com/masashitsubaki/CPI_prediction/tree/master/. We used 3D structures of proteins in Human dataset from https://github.com/prokia/drugVQA. Also, all instructions and codes for our experiments are available at https://github.com/yazdanimehdi/AttentionSiteDTI

## Acknowledgments

We thank Ms.Katalina Biondi for discussions and her valuable feedback and comments on earlier versions of the manuscript.

## Biographical Note

**Mehdi Yazdan-Jahromi** is a second year PhD student at University of Central Florida. His current research interests include Graph Neural Networks, Algorithmic Fairness.

**Niloofar Yousefi**PhD (University of Central Florida) is a Postdoctoral Research Associate with the emphasize on Computer Science, Machine Learning and Agent-based Modeling at UCF’s Complex Adaptive Systems (CAS) laboratory in the collage of Engineering and Computer Science. **Aida Tayebi** is a second year PhD student at University of Central Florida. Her current research interests include Algorithmic Fairness, and bias mitigation techniques in DTI. **Ozlem Ozmen Garibay** is an Assistant Professor of Industrial Engineering and Management System at the University of Central Florida where she directs the Human-Centered Artificial Intelligence Research Lab (Human-CAIR Lab). Prior to that, she served as the Director of Research Technology. Her areas of research are big data, social media analysis, social cybersecurity, artificial social intelligence, human-machine teams, social and economic networks, network science, STEM education analytics, higher education economic impact and engagement, artificial intelligence, evolutionary computation, and complex systems.

**Sudipta Seal** is currently the chair of the Department of Materials Science and Engineering at University of Central Florida, as well as a Pegasus Professor and a University Distinguished Professor. He joined the Advanced Materials Processing and Analysis Center and UCF in 1997. He has been consistently productive in research, instruction and service to UCF since 1998. He has served as the Nano Initiative coordinator for the vice president of research and commercialization. He served as the director of AMPAC and the NanoScience Technology Center from 2009 to 2017.

**Elayaraja Kolanthai** Ph.D. (Anna University) is a Postdoctoral Research Associate at UCF’s Materials Science and Engineering. His current research interests include Development of nanoparticles, layer-by-layer antimicrobial/antiviral nanoparticle coating, polymer composites for tissue engineering, and gene/drug delivery application.

**Craig J. Neal**PhD (University of Central Florida) is a Postdoctoral Research Associate at UCF’s Materials Science and Engineering. His current research interests include Wet chemical synthesis and surface engineering of nanoparticles for biomedical applications and electrochemical devices. Electroanalysis of nanomaterials and bio-nano interactions.

